# Cloud-Connected Pluripotent Stem Cell Platform Enhances Scientific Identity in Underrepresented Students

**DOI:** 10.64898/2026.01.19.699793

**Authors:** Samira Vera-Choqqueccota, Drew Ehrlich, Vladimir Luna-Gomez, Sebastian Hernandez, Jesus Gonzalez-Ferrer, Hunter E. Schweiger, Kateryna Voitiuk, Yohei Rosen, Kivilcim Doganyigit, Isabel Cline, Rebecca Ward, Erika Yeh, Karen H. Miga, Barbara Des Rochers, Sri Kurniawan, David Haussler, Kristian López Vargas, Mircea Teodorescu, Mohammed A. Mostajo-Radji

## Abstract

Stem cell research offers unique opportunities for authentic scientific engagement, yet infrastructure requirements have confined participation to elite institutions, perpetuating workforce disparities. We developed an integrated framework combining engineered biology, cloud-connected microscopy, and validated psychometric assessment to make pluripotent stem cell (PSC) experimentation widely accessible. The framework comprises three components: a doxycycline-inducible *NGN2* mouse embryonic stem cell line for rapid neuronal specification, low-cost cloud microscopy for remote observation, and the validated Stem Cell Research Identity Scale (SCRIS) for quantifying educational outcomes. Implementation across a Title I high school and urban community college demonstrated significant increases in scientific identity. Students using differentiating PSCs showed broader science identity development than those using neuroblastoma cells, particularly in competence, research readiness, and recognition. High school students showed enhanced research competence gains compared to community college students despite equivalent intervention duration. Demographic analyses revealed enhanced effectiveness for Hispanic and first-generation college students. This framework provides a scalable model for broadening participation in advanced biomedical research.

## INTRODUCTION

Pluripotent stem cell (PSC) technologies have redefined our capacity to model human development and disease, yet the pipeline for research participation remains geographically and socioeconomically constrained [1, 2]. While regenerative medicine promises to revolutionize global health, the specialized infrastructure required for PSC research, including stringent biosafety protocols, has confined the field to well-resourced institutions [3–6]. This centralization creates persistent participation gaps that disproportionately exclude students and researchers from community colleges, minority-serving institutions, and under-resourced secondary schools [7, 8]. Despite initiatives such as the California Institute for Regenerative Medicine (CIRM) prioritizing workforce development, translating cutting-edge research into accessible educational experiences remains a fundamental challenge [9]. Preparing a diverse workforce for the rapidly evolving biomedical landscape requires addressing both pedagogical and infrastructure barriers simultaneously.

Project-based learning (PBL) has demonstrated effectiveness in enhancing engagement and scientific reasoning by allowing students to pursue authentic, hypothesis-driven investigations [10, 11]. This approach improves learning outcomes in biotechnology by promoting autonomy and stronger connections to scientific practice [12–16]. PBL shows particularly strong benefits for students from underrepresented backgrounds [15, 17–19]. A critical outcome of such pedagogical models is the development of scientific identity, defined as the extent to which students see themselves as scientists and envision scientific careers [20, 21]. Scientific identity encompasses dimensions of competence, performance, recognition, and interest, all of which predict persistence in STEM fields [22]. While validated instruments exist to measure science identity broadly [23], specialized fields like stem cell biology lack tools that capture their unique conceptual frameworks and technical trajectories. This absence limits systematic evaluation of educational interventions in regenerative medicine.

Cloud-enabled tissue culture technologies provide a solution by enabling remote access and real-time monitoring through integrated imaging and data acquisition platforms [4, 24, 25]. Previous implementations of cloud-connected microscopy have improved learning outcomes in diverse classrooms but have relied primarily on transformed cell lines, such as neuroblastoma cells and other simple models, which lack the developmental relevance of PSC systems [24–31]. Extending cloud-based experimentation to authentic stem cell biology requires overcoming timeline constraints, as differentiation protocols typically require weeks of meticulous maintenance and validated developmental endpoints. These requirements have confined stem cell research to specialized laboratory environments with dedicated personnel.

Here, we present an integrated framework addressing biological, technological, and assessment barriers to stem cell education simultaneously. We engineered a mouse embryonic stem cell (mESC) line with doxycycline-inducible *NGN2* that enables rapid cortical neuronal specification within five days, compatible with standard academic timelines. We deployed a low-cost cloud-connected microscopy platform for distributed real-time observation. To quantify educational impact, we developed and validated the Stem Cell Research Identity Scale (SCRIS), a psychometrically robust instrument measuring four dimensions of scientific identity in regenerative medicine contexts. Implementation across a Title I high school and an urban community college demonstrated that authentic stem cell research can be effectively replicated outside traditional laboratory environments with measurable science identity gains.

This work provides a validated blueprint for making regenerative medicine education widely accessible and establishes rigorous methodology for quantifying the impact of accessible scientific infrastructure.

## RESULTS

### A Cloud-Integrated Platform Enables Rapid Neuronal Specification

To enable stem cell research within constrained academic timelines, we engineered a mESC line with doxycycline-inducible expression of the proneural transcription factor *NGN2* (Figure 1). The genetic architecture comprises a human-codon optimized *NGN2* transgene under TetO control integrated via PiggyBac transposition, alongside a constitutively expressed TagBFP reporter and puromycin resistance gene (Figure 1A). Following antibiotic selection, fluorescence-activated cell sorting enriched for high and uniform BFP expression, ensuring homogeneous starting populations for distributed experiments (Figure S1). Quantitative PCR using human codon-specific primers confirmed robust transgenic *NGN2* induction following doxycycline administration, with 2 and 4 *µ*g/mL producing comparable maximal expression (Figure 1B).

**Figure 1.**
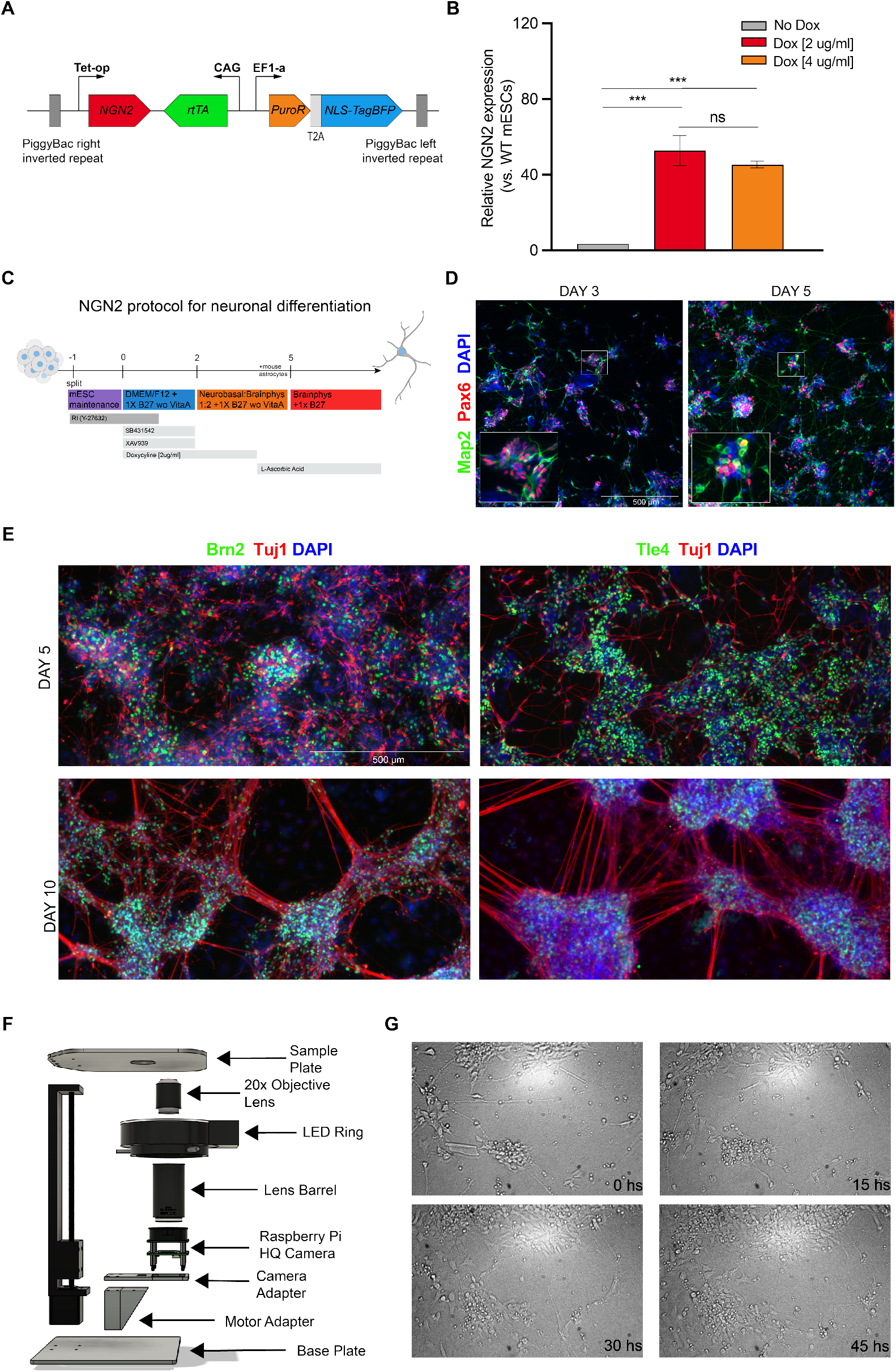
Engineering and validation of a doxycycline-inducible *NGN2* mESC line and neuronal differentiation workflow. **(A)** Schematic representation of the genetic cassette encoding a doxycycline-inducible *NGN2* transgene under TetO control, together with rtTA, and a constitutive EF1*α*-driven PuroR-NLS-TagBFP selection marker. **(B)** Quantitative PCR (qPCR) analysis of *NGN2* expression relative to wild-type (WT) mESCs under no doxycycline and increasing doxycycline concentrations (2 and 4 *µ*g/mL). Doxycycline treatment induces robust upregulation of *NGN2*. Data are presented as mean ± SEM; \*\*\**p* < 0.001; ns, not significant (unpaired t-test). **(C)** Schematic overview of the culture media composition and timeline used for neuronal differentiation following *NGN2* induction, including transitions from mESC maintenance medium to neural induction and BrainPhys-based maturation media. **(D)** Immunofluorescence analysis at day 3 and day 5 showing expression of early neural (Pax6) and neuronal markers (Map2), with nuclei counterstained with DAPI. **(E)** Immunofluorescence characterization of neuronal subtype-associated markers Brn2 and Tle4 at day 5 and day 10, co-stained with the panneuronal marker Tuj1 and DAPI, demonstrating progressive neuronal maturation over time. Scale bars, 500 *µ*m. **(F)** Schematic of the modular imaging system, including a 20× objective, LED illumination, Raspberry Pi HQ camera, and motorized base. **(G)** Representative time-lapse images of *NGN2*-induced neurons derived from stem cells starting at day 5 post-differentiation, shown over a 45-hour imaging period.

We optimized a differentiation protocol using Wnt inhibition via XAV939 and TGF-*β* inhibition via SB431542 to promote rapid neural commitment in a 2D format (Figure 1C) [32]. While cells in maintenance media showed no neural features, the combination of small molecule inhibitors and *NGN2* induction produced synchronous differentiation within 48 hours of doxycycline administration (Figure S2). Immunofluorescence analysis revealed progressive neuronal differentiation. By day 3, cultures showed extensive Pax6 expression marking cortical neural progenitors, with initial Map2-positive neurons emerging (Figure 1D). By day 5, Map2-positive neuronal populations expanded substantially (Figure 1D). To assess cortical identity, we examined expression of Brn2, which labels intermediate progenitors and callosal projection neurons (CPNs), and Tle4, which marks corticofugal projection neurons (CFuPNs) (Figure 1E). Both markers were robustly expressed by day 5, confirming acquisition of cortical layer-specific identities. By day 10, Tuj1 staining revealed extensive axodendritic processes characteristic of maturing cortical neurons (Figure 1E). This compressed timeline enables complete observation of pluripotency-to-neuron transitions and cortical specification within five days, compatible with standard academic schedules while maintaining developmental authenticity.

To enable remote observation across bandwidth-limited environments, we developed a low-cost cloud-connected microscopy platform optimized for continuous operation within standard tissue culture incubators (Figure 1F-G). The system achieves 2 to 3 *µ*m resolution at 20*×* magnification with automated image capture at 90 to 120 second intervals. To ensure accessibility across varying network conditions, we implemented cloud streaming via YouTube’s BitRate streaming infrastructure [33]. This pragmatic choice dynamically adjusts video quality in response to fluctuating bandwidth, allowing students in resource-constrained settings to maintain continuous observation without requiring institutional network upgrades. By leveraging existing commercial streaming infrastructure, we minimize technical barriers to adoption while ensuring high-fidelity biological observation.

### The SCRIS Provides Validated Measurement of Scientific Identity in Stem Cell Education

To rigorously assess how participation in stem cell research influences scientific identity, we developed the SCRIS, a multidimensional instrument grounded in established science identity frameworks and adapted for the unique demands of regenerative medicine education [20, 21]. The design process was informed by theoretical and empirical studies on science identity, with five initial dimensions identified: performance, competence, recognition, interest, and professional role models, comprising a total of 24 items (Table S1). Items were adapted from existing validated instruments and modified to reflect stem cell research context. [34–37].

The first validation step involved exploratory factor analysis (EFA) using data from 133 students enrolled in the University of California, Santa Cruz course BME80G: Bioethics in the 21st Century: Science, Business, and Society (Table S2). This population was selected for its disciplinary diversity and exposure to stem cell concepts without specialized training. EFA was conducted using principal axis factoring with Promax rotation and Kaiser normalization. Bartlett’s test of sphericity was significant (*χ*^2^ = 3540.659, *p* < 0.001), and the Kaiser-Meyer-Olkin measure was 0.890, exceeding recommended thresholds [38]. Analysis revealed a four-factor model explaining 73.9% of total variance (Figure 2A). Factor 1 (3.9% of the variance) was defined as Performance and Self-Efficacy. Factor 2 (18.7% of the variance) corresponded to Competence and Research Readiness. Factor 3 (8.2% of the variance) represented Recognition. Factor 4 (43.1% of the variance) combined interest and role model items and was designated Career Interest and Role Models.

**Figure 2.**
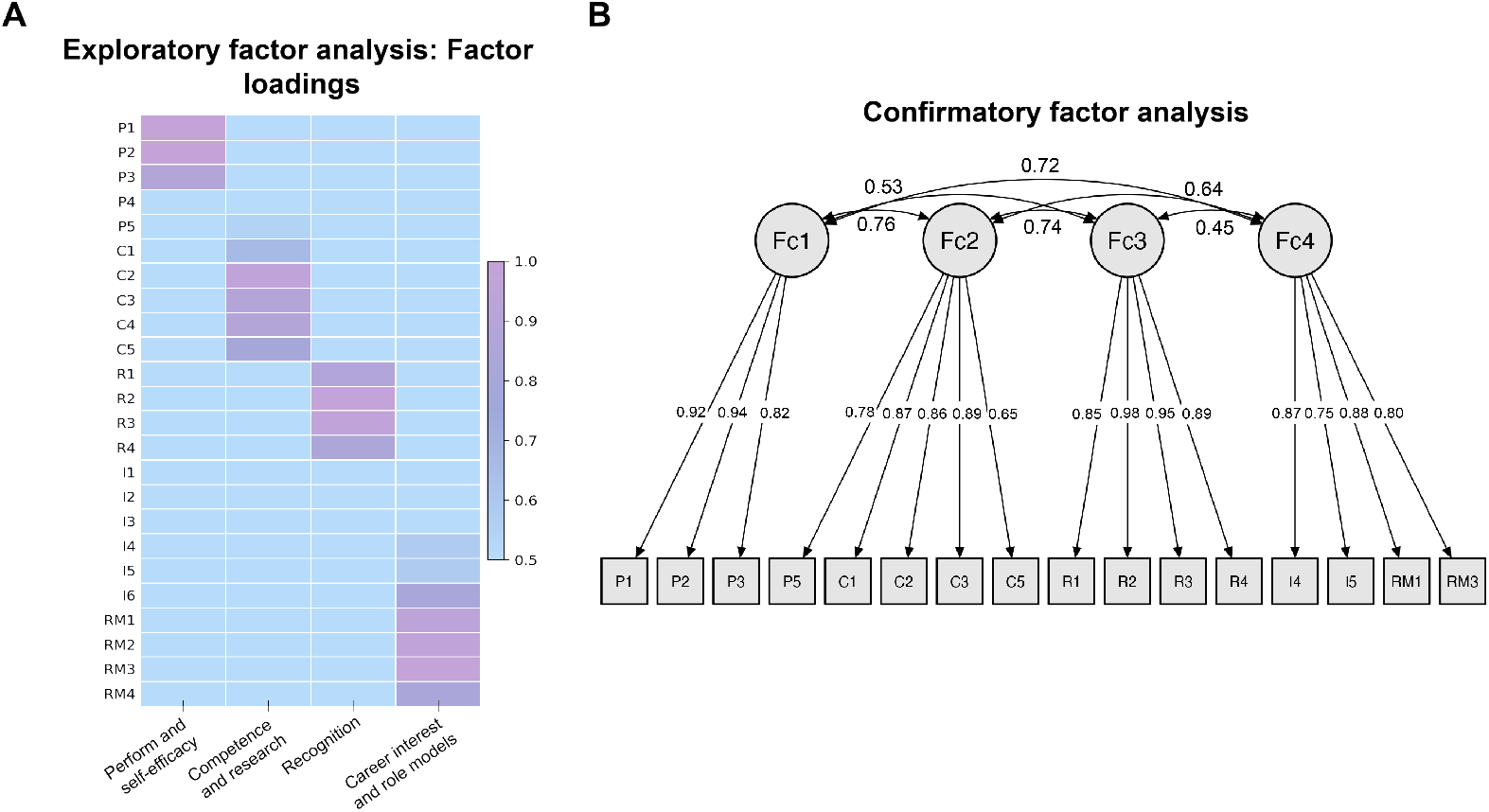
Validation of the Stem Cell Research Identity Scale (SCRIS). (**A**) Exploratory factor analysis (EFA) of UCSC undergraduate survey responses (N = 133). (**B**) Confirmatory factor analysis (CFA) of students from Alisal High School and Berkeley City College (N = 95) confirmed the four-factor model with strong overall model fit.

To confirm this structure in our target populations, we performed confirmatory factor analysis (CFA) on 95 students from diverse educational settings: AP Biology classes at Alisal High School, a Title I school serving a predominantly agricultural community in Salinas, CA, and the Biotechnology Program at Berkeley City College, an urban community college with significant populations of adult learners and first-generation immigrants (Figure 2B), (Tables S3-S7). Initial model evaluation indicated that the removal of select items with low factor loadings or cross-loadings was necessary to improve overall model fit. Following this refinement process, the final CFA model retained 16 items distributed across four factors (Table S8). The refined model demonstrated strong fit across multiple indices (CMIN/DF = 1.324, TLI = 0.97, CFI = 0.975, RMSEA = 0.058, SRMR = 0.059). All standardized factor loadings exceeded 0.50 and were statistically significant (*p* < 0.001) [39].

Cronbach’s alpha coefficients indicated strong internal consistency for each factor: Performance and Self-Efficacy (*α* = 0.903), Competence and Research Readiness (*α* = 0.888), Recognition (*α* = 0.954), and Career Interest and Role Models (*α* = 0.888), with high overall instrument reliability (*α* = 0.940) [40]. The final SCRIS thus comprises 16 items, providing a concise and statistically robust instrument for assessing scientific identity formation in stem cell education contexts.

### Five-Phase Framework Delivers Authentic Stem Cell Research Across Distributed Sites

We implemented a five-phase PBL framework across Alisal High School and Berkeley City College, both utilizing the *NGN2* differentiation platform (Figure 3A). The curriculum included: (1) instructional preparation via Zoom covering stem cell biology fundamentals and pre-intervention SCRIS administration, (2) student-driven experimental design with culturally relevant compound selection, (3) collaborative execution with real-time cloud microscopy access via Zoom connection and YouTube streaming supported by a cloud-connected experimental platform (Figure 3B), (4) open-ended data analysis emphasizing student-defined metrics, and (5) peer presentation, reporting, and post-intervention SCRIS assessment. Students proposed testing compounds ranging from traditional remedies suggested by family members to environmental toxins of local concern, fostering personal investment in the research questions.

**Figure 3.**
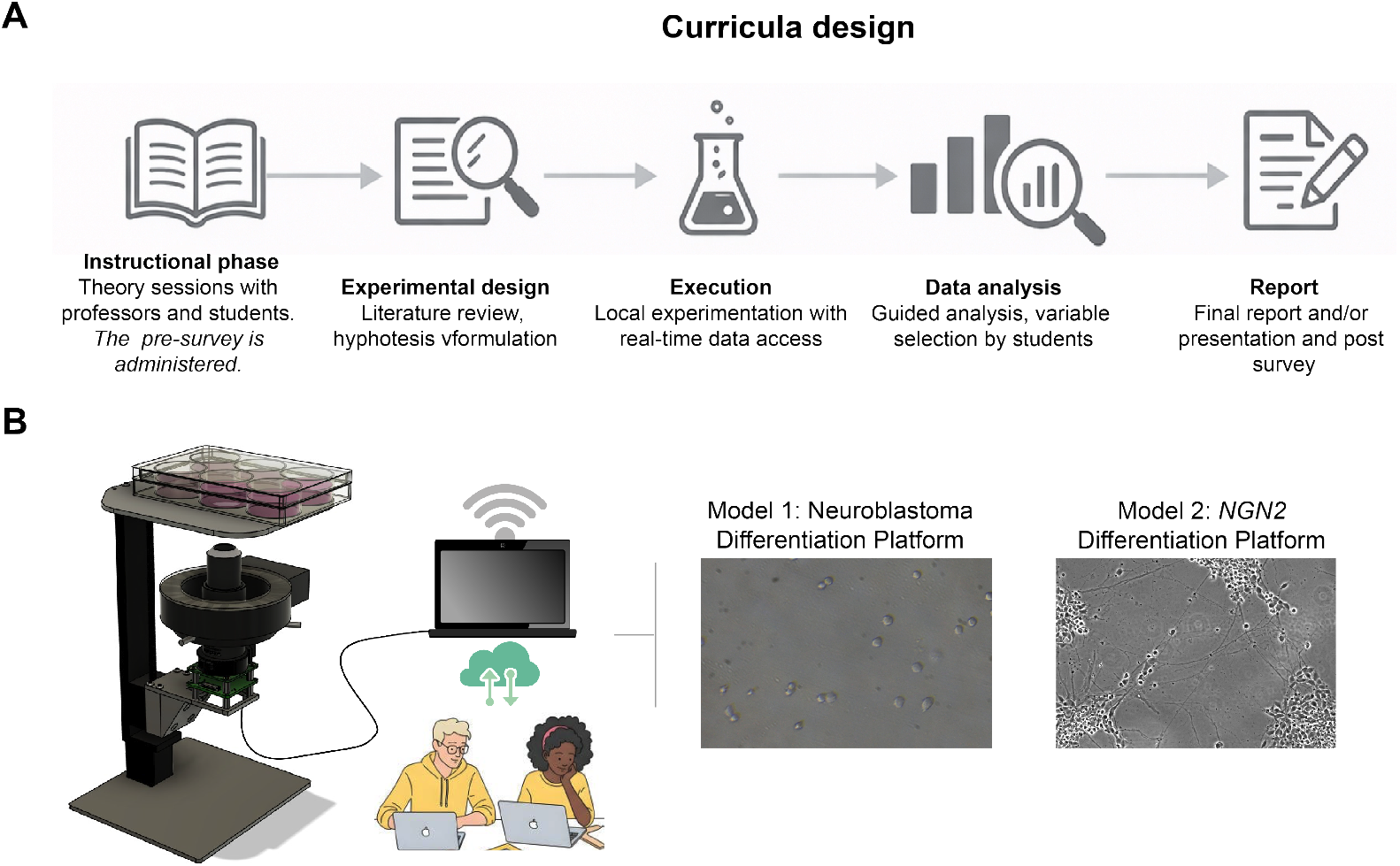
Five-phase remote experimental framework integrating curriculum and cloud-based experimentation. (**A**) Schematic of the five-phase remote PBL workflow: instructional preparation, experimental design, experimental execution, data analysis, and reporting, including pre- and postintervention assessments. (**B**) Overview of the remote experimental platform combining a custom-built, cloud-connected microscope with live data streaming, enabling students to access and analyze experiments conducted locally. Neuroblastoma and *NGN2* differentiation platforms were used as biological models.

### Pluripotent Stem Cells Enhance Research Identity Compared to Neuroblastoma Cells

To examine whether starting cell type influences science identity development, we compared SCRIS outcomes between students using PSCs versus neuroblastoma cells. Building on previous work demonstrating that remote PBL with SH-SY5Y neuroblastoma differentiation enhances STEM identity [24], we collected SCRIS data from a neuroblastoma differentiation platform cohort at Alisal High School (2024, N = 21) and compared outcomes to the *NGN2* differentiation platform cohort (2025, N = 32). Both cohorts were taught by the same instructor using comparable cloud-based project frameworks, allowing exploration of biological system effects while acknowledging temporal separation. Pre-intervention SCRIS scores were statistically equivalent between cohorts across all four dimensions (Figure 4A, all *p* > 0.05), indicating comparable baseline populations.

**Figure 4.**
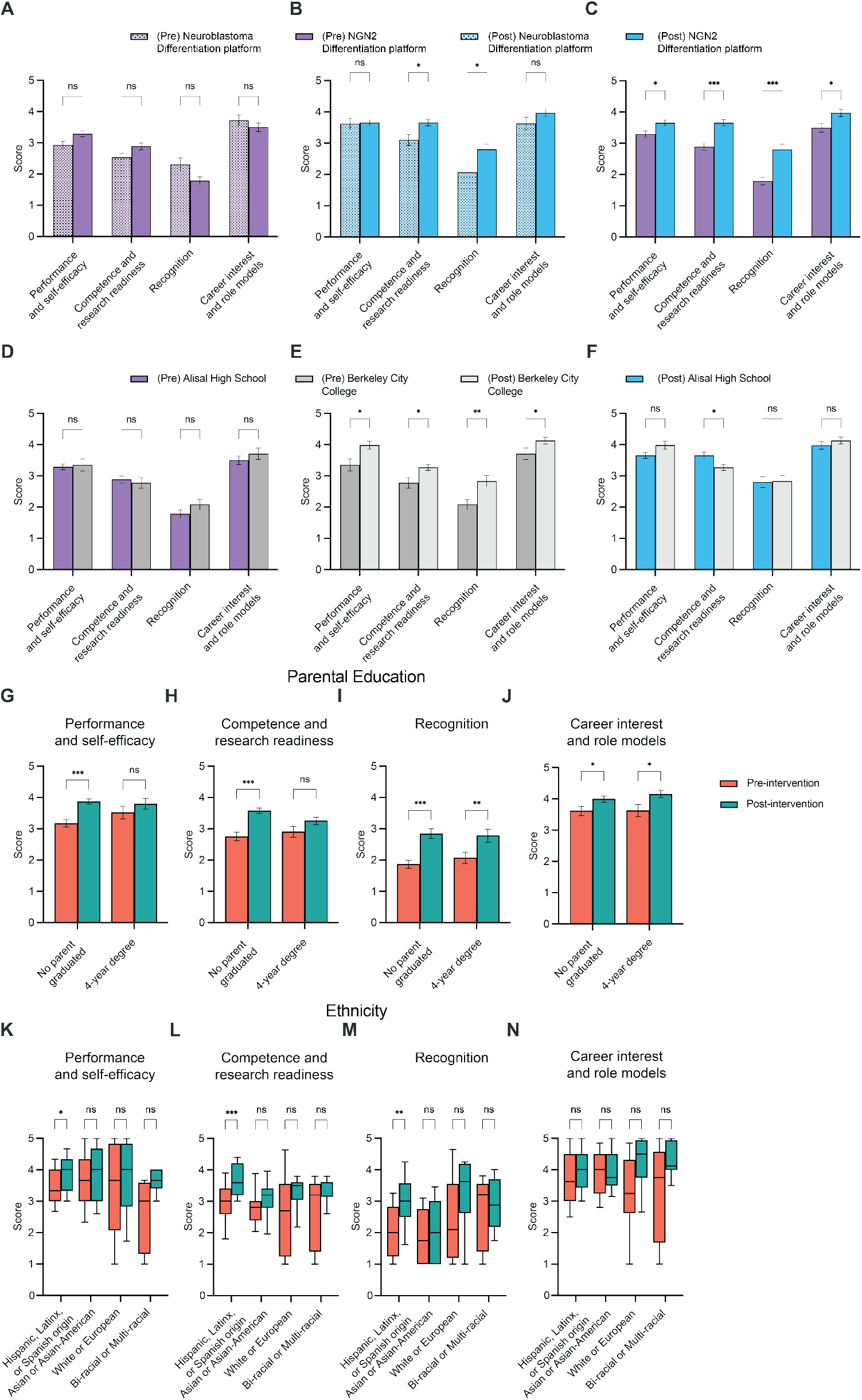
Science identity formation outcomes across biological systems and institutional contexts. (**A**) Pre-intervention comparison between neuroblastoma differentiation platform (2024) and *NGN2* differentiation platform (2025) cohorts at Alisal High School shows equivalent baseline identity across all dimensions. (**B**) Post-intervention scores between neuroblastoma differentiation platform (2024) and *NGN2* differentiation platform (2025) cohorts. (**C**) Alisal High SchooltextitNGN2 differentiation platform (2025) cohort demonstrates significant pre-to-post gains across all four SCRIS dimensions. (**D**) Pre-intervention comparison between Alisal High School and Berkeley City College, both using textit-NGN2 differentiation platform, shows equivalent baseline identity across all dimensions. (**E**) Berkeley City College textitNGN2 differentiation platform cohort demonstrates significant pre-to-post gains across all four SCRIS dimensions. (**F**) Post-intervention comparison between Alisal High School and Berkeley City College. (**G–J**) Stratified analysis by parental educational attainment (no college degree vs. ≥4-year degree) using aggregated *NGN2* cohorts from Alisal High School and Berkeley City College. Bar plots represent mean ± SEM; group differences were assessed using unpaired two-tailed t-tests. (**K–N**) Stratified analysis by self-reported ethnicity (Hispanic/Latinx, Asian or Asian-American, White or European, and Bi-racial or Multiracial) using aggregated *NGN2* cohorts from Alisal High School and Berkeley City College. Boxplots display median and interquartile range; group differences were assessed using non-parametric Wilcoxon rank-sum tests. Sample sizes: neuroblastoma differentiation platform cohort (N = 21), Alisal *NGN2* differentiation platform cohort (N = 32), Berkeley *NGN2* differentiation platform cohort (N = 42); parental education groups: no college degree (N = 42), ≥4-year degree (N = 32); ethnicity groups: Hispanic/Latinx (N = 34), Asian or Asian-American (N = 15), White or European (N = 12), Bi-racial or Multiracial (N = 8). \*\*\**p* < 0.001, \*\**p* < 0.01, \**p* < 0.05; ns, not significant.

Post-intervention analysis revealed distinct patterns of science identity development between the two biological systems (Figure 4B). While both cohorts showed significantly higher post-intervention scores compared to pre-intervention scores in Performance and Self-Efficacy and Competence and Research Readiness the *NGN2* differentiation platform produced a broader enhancement across science identity dimensions. Most notably, students using the *NGN2* differentiation platform cohort demonstrated significantly higher post-intervention scores in Recognition (*p* < 0.001) and Career Interest and Role Models (*p* < 0.05), whereas the neuroblastoma differentiation cohort showed minimal differences between pre- and post-intervention scores in these dimensions. Recognition and Career Interest and Role Models post-intervention scores were significantly higher than pre-intervention scores in the *NGN2* differentiation platform cohort (*p* < 0.001 and *p* < 0.05) but did not differ significantly in the neuroblastoma cohort (Figure S3). These findings suggest that authentic developmental systems with lineage specification and technical complexity engage students more deeply across multiple facets of scientific identity, particularly in dimensions related to research capability, professional recognition, and career aspiration.

### Platform Produces Differential Science Identity Gains Across Institutional Contexts

To assess whether educational setting influences outcomes, we compared the Alisal High School (AP Biology, N = 32) and Berkeley City College (Biotechnology Program, N = 42) cohorts, both using the *NGN2* stem cell platform with identical intervention duration and structure. Pre-intervention science identity was statistically equivalent across all SCRIS dimensions (Figure 4D, all *p* > 0.05), indicating comparable baseline populations despite different institutional contexts. Both cohorts showed significantly higher post-intervention scores compared to pre-intervention scores across all four identity dimensions (Figures 4C, E).

Post-intervention analysis revealed institutional differences in one specific dimension (Figure 4F). Alisal High School students showed significantly higher Competence and Research Readiness compared to Berkeley City College students (*p* < 0.05). This finding likely reflects baseline differences in academic preparation. Alisal students were enrolled in AP Biology, a course that typically attracts academically focused students with recent biology coursework. Berkeley students were enrolled in Introductory Biotechnology, the first course in a three-semester program, with many adult learners who had not taken biology in years. Both groups entered with equivalent baseline science identity scores (Figure 4D), but the AP Biology students’ stronger academic foundation may have facilitated larger competence gains from the same intervention. Performance and Self-Efficacy, Recognition, and Career Interest and Role Models were equivalent between institutions (all *p* > 0.05).

### First-Generation and Hispanic Students Show Enhanced Science Identity Development

To understand whether science identity changes varied across student demographics, we conducted independent t-tests comparing pre-intervention to post-intervention scores within each demographic sub-group.

First-generation college students, defined as those whose parents did not complete bachelor’s degrees, showed significant increases across all science identity dimensions Figure 4G-J). Performance and Self-Efficacy increased significantly from pre-to post-intervention (Figure 4G), *p* < 0.001), as did Competence and Research Readiness (Figure 4H, *p* < 0.001) and Recognition (Figure 4I, *p* < 0.001). Career Interest and Role Models increased significantly in both first-generation students (*p* < 0.05) and continuing-generation students whose parents completed four-year degrees (*p* < 0.05) (Figure 4J). Notably, continuing-generation students showed significant gains only in Recognition (Figure 4I, *p* < 0.01) and Career Interest (Figure 4J), with no significant changes in Performance and Self-Efficacy or Competence and Research Readness (both *p* > 0.05).

Hispanic students demonstrated significant increases in Performance and Self-Efficacy (Figure 4K, *p* < 0.05), Competence and Research Readiness (Figure 4L, p< 0.001), and Recognition (Figure 4M, < 0.01). In contrast, students identifying as Asian or Asian-American, White or European, and Bi-racial or Multi-racial showed no significant pre-to-post changes in any dimension (Figure 4K-N, all *p* > 0.05). Career Interest and Role Models did not change significantly in any ethnic group (Figure 4N).

Analysis of gender and English language proficiency revealed that both groups within each category showed significant increases across most identity dimensions, indicating broad platform effectiveness (Figure S4). These findings suggest that authentic stem cell research delivered through accessible infrastructure produces particularly strong identity gains for first-generation college students and Hispanic students, populations historically underrepresented in STEM fields.

## DISCUSSION

We present an integrated framework addressing biological, technological, and measurement barriers to authentic stem cell education. The engineered *NGN2*-inducible mESC line enables five-day observation of pluripotency-to-neuron transitions with cortical specification (Brn2, Tle4), addressing the incompatibility between standard multi-week protocols and academic calendars while reducing instructor burden. The cloud-connected microscopy platform removes infrastructure barriers through reduced hardware costs and adaptive streaming that ensures accessibility across varying bandwidth conditions without requiring institutional network upgrades [25, 33]. The distributed observation model enables multiple students to access a single microscope simultaneously, reducing marginal costs to a few dollars per student at scale. Beyond education, the platform has potential applications in distributed research collaborations and clinical-scale manufacturing quality monitoring [41, 42]. The SCRIS provides validated measurement with strong psychometric properties, filling a gap in regenerative medicine education assessment and enabling systematic evaluation across diverse settings from high school AP Biology to community college biotechnology programs. Our findings reveal that biological authenticity, institutional context, and demographic factors influence science identity development in distinct ways. The *NGN2* differentiation platform produced substantially enhanced outcomes in Competence and Research Readiness and Recognition compared to neuroblastoma cells, dimensions directly related to students’ perception of their research capability and emerging scientist identity. Both high school AP Biology and community college introductory biotechnology students showed significant science identity gains across all dimensions, demonstrating that platform effectiveness derives from authentic biological systems and accessible infrastructure rather than requiring advanced coursework or specialized facilities.

Several considerations inform future directions. The neuroblastoma comparison involves year-over-year cohorts; concurrent randomization would strengthen causal claims. The higher competence gains observed in AP Biology students compared to introductory biotechnology students likely reflect differences in baseline academic preparation rather than institutional resources. The demographic analysis (Figure 4G-N) shows that Hispanic and first-generation students exhibited the most robust science identity and competence gains, indicating that well-designed accessible platforms can particularly benefit underrepresented populations. Qualitative and mixed-methods approaches would clarify whether differences reflect capability development or demographic-specific science identity formation. Demographic findings require replication across additional sites and larger cohorts. Longitudinal studies tracking enrollment in advanced courses and research opportunities would assess sustained career impact. Future implementations could integrate gamified virtual laboratory training [43] as preparatory experiences before cloud-connected experiments. Extension to human induced PSCs and 3D organoid models [16, 28] would enhance translational relevance while requiring attention to complexity and cost.

These results align with workforce development priorities in regenerative medicine and initiatives to broaden STEM participation [4, 25]. The framework’s modular design enables adoption across diverse contexts, with hardware specifications, protocols, and validated SCRIS openly available. Beyond education, the cloud infrastructure demonstrates proof of principle for distributed biological research, multi-site collaborations, clinical manufacturing quality monitoring, and international partnerships [41, 42, 44]. By integrating authentic developmental biology, autonomous observation systems, and validated psychometric assessment into a scalable framework, this work provides practical tools and empirical evidence that cutting-edge biomedical research can be made widely accessible without compromising scientific rigor or educational outcomes.

## RESOURCE AVAILABILITY

### Lead contact

Further information and requests for resources should be directed to and will be fulfilled by the lead contact, Mohammed A. Mostajo-Radji.

### Materials availability

No new materials or reagents were generated in the study.

### Data and code availability

No new code was generated in this study. Anonymized individual survey data are available from the corresponding author upon reasonable request with appropriate ethical approval.

## ACKNOWLEDGMENTS

We thank the students who participated in this study. This work was supported by Schmidt Futures SF857 (D.H. and M.T.), the National Human Genome Research Institute under award RM1HG011543 (D.H. and M.T.), the National Science Foundation under awards 2134955 (D.H. and M.T.), 2034037 (M.T.), and 2515389 (D.H., M.T., and M.A.M.-R.), the CIRM awards EDUC2-12738 (B.D.R.), DISC4-16285 (M.T. and M.A.M.-R.) and DISC4-16337 (M.A.M.-R.), the University of California Office of the President award M25PR9045 (M.T. and M.A.M.-R.), the National Institute of Mental Health award U24MH132628 (M.A.M.-R. and D.H.), National Institute of Neurological Disorders and Stroke award U24NS146314 (M.A.M.-R. and D.H.), the Brain and Behavior Research Foundation 33184 (M.A.M.-R.), and the University of California Santa Cruz Center for Information Technology Research in the Interest of Society and the Banatao Institute Interdisciplinary and Innovation Program (K.L.-V. and M.A.M.-R.). H.S. was partially supported by the NSF Graduate Research Fellowship Program. S.H. received support from the UC Doctoral Diversity Initiative (DDI-UCSC-IBSC). S.V.-C. was partially supported by the Graduate Pedagogy Fellowship from the University of California Santa Cruz Teaching and Learning Center. J.G.-F. was partially supported by the QB3 Santa Cruz Graduate Innovators Fellowship. V.L.-G. was supported by the UCSC Genomics Institute Research Mentoring Internship program. Y.R. was supported by CIRM postdoctoral fellowships under Award number EDUC4-12759 and the Institute for the Biology of Stem Cells (IBSC) at UC Santa Cruz and by the Peggy and Jack Baskin fellowship and UCSC Baskin School of Engineering fellowship programs. We also thank the UC Santa Cruz Life Sciences Microscopy Center Core Facility (RRID:SCR_021135) and the UCSC Flow Cytometry Core (RRID:SCR_021149) supported by CIRM Major Facility Award (FA1-00617 to UCSC).The content is solely the responsibility of the authors and does not necessarily represent the official views of the University of California, the National Institutes of Health, the National Science Foundation, CIRM, or any other federal or state agency.

## AUTHOR CONTRIBUTIONS

S.V.-C. and M.A.M.-R. conceived the experiments. S.V.-C., D.E., V.L-G., S.H., J.G.-F., H.E.S., K.V. and Y.R. performed the experiments. K.D., I.C., R.W., E.Y., K.M., and D.H. led the courses where these experiments were performed. B.D.R., S.K., K.L.-V., D.H., M.T. and M.A.M.-R. supervised the work. K.L.-V., D.H., M.T. and M.A.M.-R. secured the funding.

## DECLARATION OF GENERATIVE AI AND AI-ASSISTED TECHNOLO-GIES

During the preparation of this work, the authors used ChatGPT and Claude to improve language clarity and readability. All content was reviewed and edited by the authors, who take full responsibility for the final version.

## MATERIALS AND METHODS

### Ethics Statement and Institutional Oversight

This study was reviewed and approved by the University of California Santa Cruz Office of Research Compliance Administration and Institutional Review Board under protocol number HS-FY2023-264.

### Engineered Mouse Embryonic Stem Cell Culture

All experiments utilized an adapted C57/BL6 mESC line derived from a male of the C57/BL6J strain (Millipore Sigma #SF-CMTI-2). Mycoplasma testing confirmed the lack of contamination. Cells were maintained on Recombinant Human Protein Vitronectin (Thermo Fisher Scientific #A14700)-coated plates. Vitronectin coating was incubated for 30 minutes at a concentration of 0.5 μg/mL dissolved in 1X Phosphate Buffered Saline pH 7.4 (Thermo Fisher Scientific #70011044). Maintenance utilized mESC media containing Glasgow Minimum Essential Medium (Thermo Fisher Scientific #11710035), Embryonic Stem Cell Qualified Fetal Bovine Serum (Thermo Fisher Scientific #10439001), 0.1 mM MEM Non-Essential Amino Acids (Thermo Fisher Scientific #11140050), 1 mM Sodium Pyruvate (Millipore Sigma #S8636), 2 mM Glutamax supplement (Thermo Fisher Scientific #35050061), 0.1 mM 2-Mercaptoethanol (Millipore Sigma #M3148), and 0.05 mg/mL Primocin (Invitrogen #ant-pm-05). Media was supplemented with 1,000 units/mL of ESGRO Recombinant Mouse Leukemia Inhibitory Factor (Millipore Sigma #ESG1107). Media was changed daily. Dissociation and passages were performed using ReLeSR passaging reagent (Stem Cell Technologies #05872). Cell freezing used mFreSR cryop-reservation medium (Stem Cell Technologies #05855).

### Genetic Engineering of Doxycycline-Inducible *NGN2* mESC Line

mESCs were co-transfected with a PiggyBac transposon vector encoding TetO-*NGN2*-T2A-rtTA and EF1*α*-PuroR-NLS-TagBFP (Addgene #198397), with a Super PiggyBac transposase–expressing plasmid provided by the Colquitt laboratory, using Lipofectamine 3000 (Thermo Fisher Scientific, #L3000015) according to the manufacturer’s instructions. The *NGN2* sequence in this vector is codon-optimized for humans. Following 48 hours recovery, puromycin selection (1 μg/mL, Invitrogen #A1113803) was applied for 5 days. BFP-positive cells were enriched via FACS on a BD FACSAria II, gating for the top 11% of fluorescence intensity.

### Quantitative PCR Analysis

Total RNA was extracted using the RNeasy Mini Kit (Qiagen, #74104) and reverse transcribed using the SuperScript III First-Strand Synthesis System (Invitrogen, #18080051). Quantitative PCR was performed on a QuantStudio 3 Real-Time PCR System (Applied Biosystems) using TaqMan™ Fast Advanced Master Mix (Thermo Fisher Scientific, #4444557). Expression levels of *NGN2* were normalized to *Gapdh* using the ΔΔC_t_ method.

### Inducible *NGN2* Neuronal Differentiation

To generate *NGN2* neurons, mESCs were dissociated using TrypLE Express Enzyme (Thermo Fisher Scientific #12604021) for 7 minutes at 37°C and seeded at a density of 300,000 cells in 2 mL of maintenance media per well in 6-well plates that were pre-coated with poly-L-ornithine (0.01%) (Millipore Sigma #P4957) and vitronectin for 1 hour, washed with PBS (1X) three times, and subsequently coated with laminin (15 *µ*g/mL) (Corning #354232) and fibronectin (5 *µ*g/mL) (Corning #354008) . Twentyfour hours prior to differentiation induction (day -1), media was supplemented with Rho Kinase Inhibitor Y-27632 (10 μM, Tocris #1254).

On day 0, differentiation was initiated by replacing the media with neural induction medium containing DMEM/F12 with GlutaMAX (Thermo Fisher Scientific #10565018), 0.1 mM MEM Non-Essential Amino Acids, 1 mM Sodium Pyruvate, and 0.05 mg/mL primocin. This medium was supplemented with 2 μg/mL doxycycline (Sigma #D9891) to induce *NGN2* expression, 10 μM Rho Kinase Inhibitor Y-27632, 5 μM XAV939 (Tocris #3748), and 10 μM TGF-*β* inhibitor SB431542 (Tocris #1614). Doxycycline, XAV939, and SB431542 were maintained from days 0 to 2, with daily media changes.

On day 2, media was replaced with neuronal differentiation media containing Neurobasal-A (Thermo Fisher Scientific #10565018), BrainPhys Neuronal Medium (Stem Cell Technologies #05790), 2 mM GlutaMAX supplement (Thermo Fisher Scientific #10565018), 1X N-2 Supplement (Thermo Fisher Scientific #17502048), 1X Chemically Defined Lipid Concentrate (Thermo Fisher Scientific #11905031), 2 μg/mL doxycycline (until day 4), and 0.05 mg/mL primocin. On day 5 and onward, cultures were transitioned to maturation media containing BrainPhys Neuronal Medium, 1X N-2 Supplement, 1X Chemically Defined Lipid Concentrate, 1X B-27 Supplement (Thermo Fisher Scientific #17504044), 0.5% (v/v) Matrigel GFR Basement Membrane Matrix (LDEV-free) (Corning #354230), 0.05 mg/mL primocin, and 5 μg/mL heparin (Sigma #H3149). Media was changed every other day from day 5 onward.

### Neuroblastoma Differentiation

SH-SY5Y neuroblastoma cells (ATCC #CCL-131) were grown in DMEM/F-12 medium with GlutaMAX supplement (Thermo Fisher Scientific #10565018) and 10% fetal bovine serum (Thermo Fisher Scientific #26140079) at 37°C. The cells were differentiated with Neurodazine (2 μM, Millipore Sigma #N6664) and Retinoic Acid (10 μM, Millipore Sigma #R2625)

### Immunofluorescence

Cultures were fixed in 4% paraformaldehyde (Electron Microscopy Sciences #15710) for 15 minutes at room temperature, washed three times with PBS, permeabilized with 0.3% Triton X-100 in PBS for 10 minutes, and blocked in 5% normal goat serum (Jackson ImmunoResearch #005-000-121) for 1 hour at room temperature. Primary antibodies were applied overnight at 4°C: rabbit anti-Pax6 (1:500, BioLegend #901301), chicken anti-Map2 (1:1000, Abcam #ab5392), mouse anti-Tuj1 (1:1000, BioLegend #801202), rabbit anti-Brn2 (1:500, Cell Signaling #12137), mouse anti-Tle4 (1:200, Santa Cruz Biotechnology #sc-365406). Following three PBS washes, secondary antibodies (Alexa Fluor 488, 568, 647-conjugated goat anti-rabbit, anti-mouse, or anti-chicken IgG, 1:1000, Invitrogen) were applied for 1 hour at room temperature. Nuclei were counterstained with DAPI (1 μg/mL, Thermo Fisher Scientific #D1306) for 5 minutes. Images were acquired using a Zeiss live-cell widefield microscope with structured illumination and a 20*×* objective. Image processing and analysis were performed using ImageJ/Fiji software.

### Design and Psychometric Validation of the SCRIS

The SCRIS was developed in two stages. The first stage identified core dimensions based on established science identity frameworks [20, 21], including performance, competence, recognition, interest, and role models. Performance items focused on confidence in laboratory tasks and research projects while competence measured self-perceived understanding of biological concepts [34]. Recognition evaluated self-perception and external recognition as members of the scientific field. Interest gauged enthusiasm and career aspirations, and role models explored the impact of mentorship [20].

The second stage involved developing 24 items adapted from validated STEM identity measures [34–37] and modified to reflect stem cell and neuroscience contexts. The questionnaire utilized a 5-point Likert scale ranging from strongly disagree to strongly agree.

EFA was conducted using SPSS version 28.0 with principal axis factoring and Promax rotation (kappa = 4). Kaiser-Meyer-Olkin measure of sampling adequacy and Bartlett’s test of sphericity assessed data suitability for factor analysis. Items with factor loadings less than 0.40 or cross-loadings exceeding 0.32 were eliminated. The EFA sample comprised 133 students enrolled in a University of California Santa Cruz bioethics course (BME80G: Bioethics in the 21st Century). Complete demographic information for this cohort is available in (Table S2).

CFA was performed using JASP statistical software (Version 0.95.4) with maximum likelihood estimation. The CFA sample included 74 students from AP Biology classes at Alisal High School (Salinas, CA) and the Biotechnology Program at Berkeley City College (Berkeley, CA) . Model fit was evaluated using multiple indices: chi-square to degrees of freedom ratio (CMIN/DF), Tucker-Lewis Index (TLI), Comparative Fit Index (CFI), Root Mean Square Error of Approximation (RMSEA), and Standardized Root Mean Square Residual (SRMR). Internal consistency was assessed using Cronbach’s alpha coefficients for each subscale and the overall instrument. The final SCRIS retained 16 items across four factors. Complete demographic information for these cohorts is available in (Tables S3-S7).

### Distributed Cloud-Connected Microscopy Architecture

Imaging data were acquired using an open hardware in-incubator microscope adapted from published designs [24, 28, 45]. The optical system utilized a Raspberry Pi High Quality Camera module (12.3 megapixels, Sony IMX477 CMOS sensor) connected to an Arducam IMX477 UVC adapter board. A 20*×* magnification LM Plan achromatic microscope objective (Boli Optics) with 8.8 mm working distance was mounted using an RMS to C-Mount adapter (Thorlabs #RMSA5) and a 50 mm C-Mount lens tube (Thorlabs #CML50). Illumination was provided by a white LED ring light positioned beneath the culture vessel. Assembly details can be found in Supplementary Note 1.

Axial positioning used a NEMA11 stepper motor-driven linear rail with 150 mm travel distance, a 6 mm leadscrew, and a Pololu Tic T825 stepper motor driver with 24 V power supply, achieving 5 μm positioning precision. Camera control utilized the OpenCV Python library (version 4.5.4) while motor positioning was managed via the Pololu ticcmd command-line utility. Captured images were displayed on an HTML webpage and live-streamed to YouTube using Open Broadcaster Software (OBS Studio version 27.2) with RTMP protocol and H.264 encoding.

The complete assembly operated continuously within a standard tissue culture incubator at 37°C and 5% CO_2_. Automated image acquisition occurred at 90 to 120 second intervals for 3 to 5 days per experiment. Optical resolution was validated using a USAF 1951 resolution test target, confirming 2 to 3 μm resolution at 20*×* magnification.

### Educational Implementation and SCRIS Administration

Implementation followed a structured five-phase project-based learning framework conducted over 2 to 3 weeks at Alisal High School (Salinas, CA; Title I designation, 95% Hispanic enrollment, AP Biology) and Berkeley City College (Berkeley, CA; urban community college, Introductory Biotechnology course, first semester of three-semester Biotechnology Program).

#### Phase 1 (Instructional Preparation

Students received remote instruction via Zoom covering stem cell biology fundamentals, pluripotency, neuronal differentiation, the *NGN2* transcription factor system, and experimental design principles. The pre-intervention SCRIS was administered electronically via Google Forms.

#### Phase 2 (Experimental Design)

Students conducted literature reviews and brainstormed experimental conditions for testing effects on differentiating neurons. Proposals included culturally relevant compounds (traditional remedies, herbs, vitamins, foods consumed locally) and environmental factors of local concern (agricultural pesticides, air pollutants, heavy metals). The class discussed proposals collectively via Zoom and selected 2-3 conditions by majority vote for parallel testing.

#### Phase 3 (Experimental Execution)

Student-selected compounds were prepared at physiologically relevant concentrations and added to parallel culture wells alongside vehicle controls. Experiments were initiated during a live Zoom session allowing students to observe cell plating, compound addition, and microscope setup. Students monitored differentiation in real-time via the YouTube live stream for 2-3 days, with phase contrast images captured automatically at 90 to 120 second intervals.

#### Phase 4 (Data Analysis)

Students downloaded time-lapse image sequences and performed analyses using self-selected metrics and freely available software (ImageJ, Excel). Analytical approaches included cell density quantification, morphological classification, neurite outgrowth measurement, cell migration tracking, and temporal dynamics of differentiation markers.

#### Phase 5 (Reporting)

Students prepared final reports or oral presentations for peers describing their experimental question, methodology, results, and biological interpretation. Post-intervention SCRIS was administered electronically.

For the neuroblastoma comparison cohort, a similar five-phase framework was implemented at Alisal High School in 2024 using SH-SY5Y neuroblastoma cells (ATCC #CCL-131), as previously described [24].

### Statistical Analysis

All statistical analyses were conducted using multiple software platforms. Exploratory factor analysis (EFA) was performed using SPSS (Version 31; IBM Corp.). Confirmatory factor analysis (CFA) was conducted using JASP statistical software (Version 0.95.4) with maximum likelihood estimation. Comparisons between pre- and post-intervention outcomes were performed using GraphPad Prism (Version 10.6.1; GraphPad Software). To maintain participant anonymity, surveys were administered without individual identifiers, preventing pairing of pre- and post-intervention responses. All comparisons therefore used independent two-tailed t-tests: comparisons of pre-intervention group means to postintervention group means within each cohort assessed intervention effects, while comparisons between cohorts at single timepoints examined baseline equivalence or outcome differences.

Demographic analyses examined whether intervention effects differed across student populations. For each demographic category, independent t-tests compared pre-intervention to post-intervention group means within each subgroup. Analyses were conducted only for demographic subgroups with sufficient sample sizes; subgroups with very small representation (e.g., one or two participants) or with no representation in one of the institutions were excluded from subgroup-specific analyses.

Normality was assessed using Shapiro–Wilk tests and visual inspection of Q–Q plots. The majority of distributions met parametric assumptions or had sufficiently large sample sizes to invoke the central limit theorem; for comparisons that violated normality assumptions, nonparametric Wilcoxon rank-sum tests were applied. Homogeneity of variance was assessed using Levene’s test. Statistical significance was set at *α* = 0.05 for all analyses. No correction for multiple comparisons was applied given the exploratory nature of demographic subgroup analyses. Data are presented as mean *±* SEM unless otherwise noted. Sample sizes are reported for each analysis.

