## Supplemental Materials for "Cloud-Connected Pluripotent Stem Cell Platform Enhances Scientific Identity in Underrepresented Students"

### 598 Supplemental Information

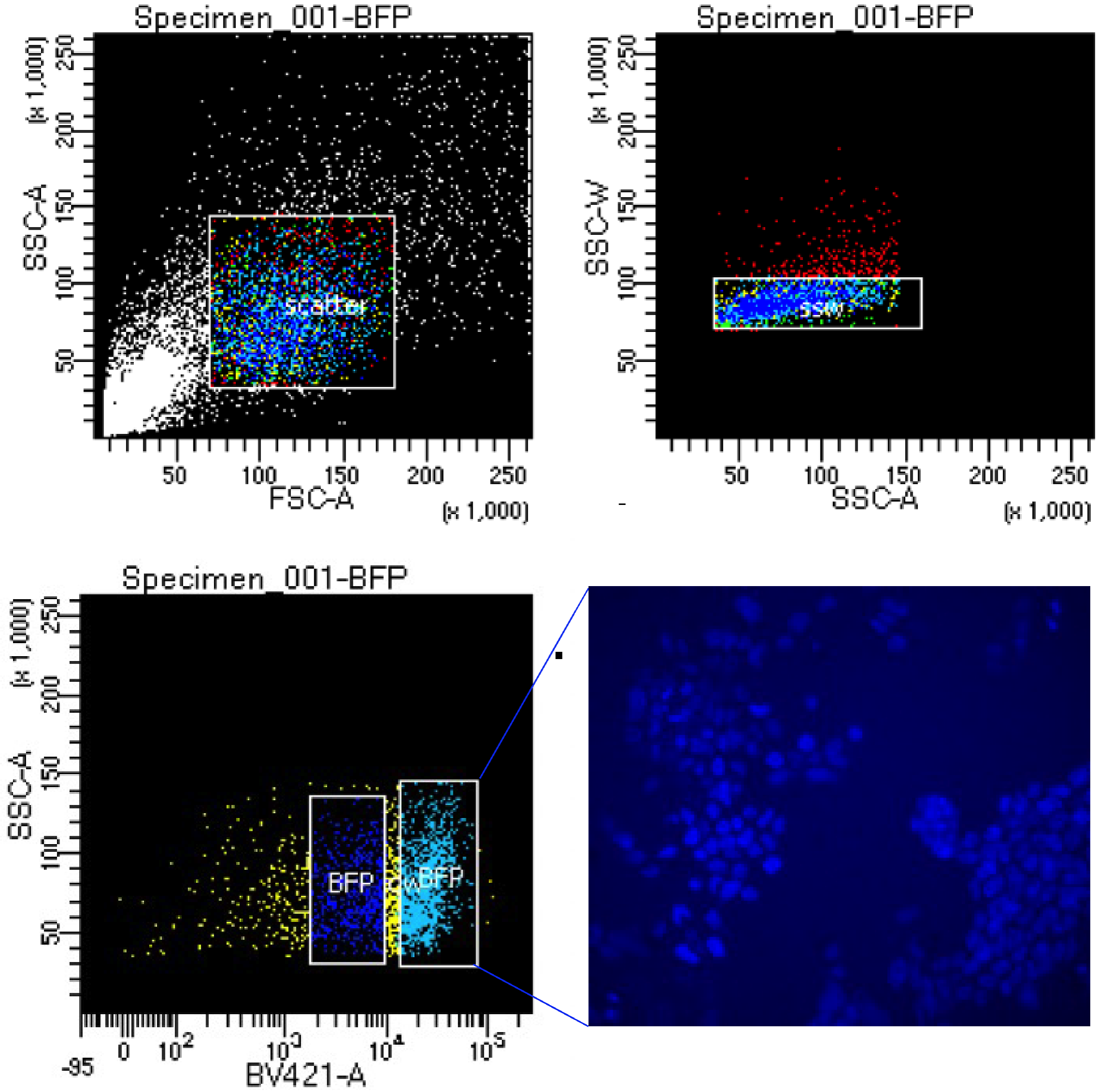


**Figure S1. FACS-based enrichment of BFP-positive cells following inducible *NGN2* cassette integration.** Related to Figure 1. Fluorescence-activated cell sorting (FACS) analysis showing the gating strategy used to enrich BFP-positive cells. Cells were gated based on FSC-A and SSC-A to exclude debris, followed by doublet exclusion and identification of BFP-positive cells based on blue fluorescence. Representative fluorescence microscopy confirms nuclear BFP expression in the sorted population.

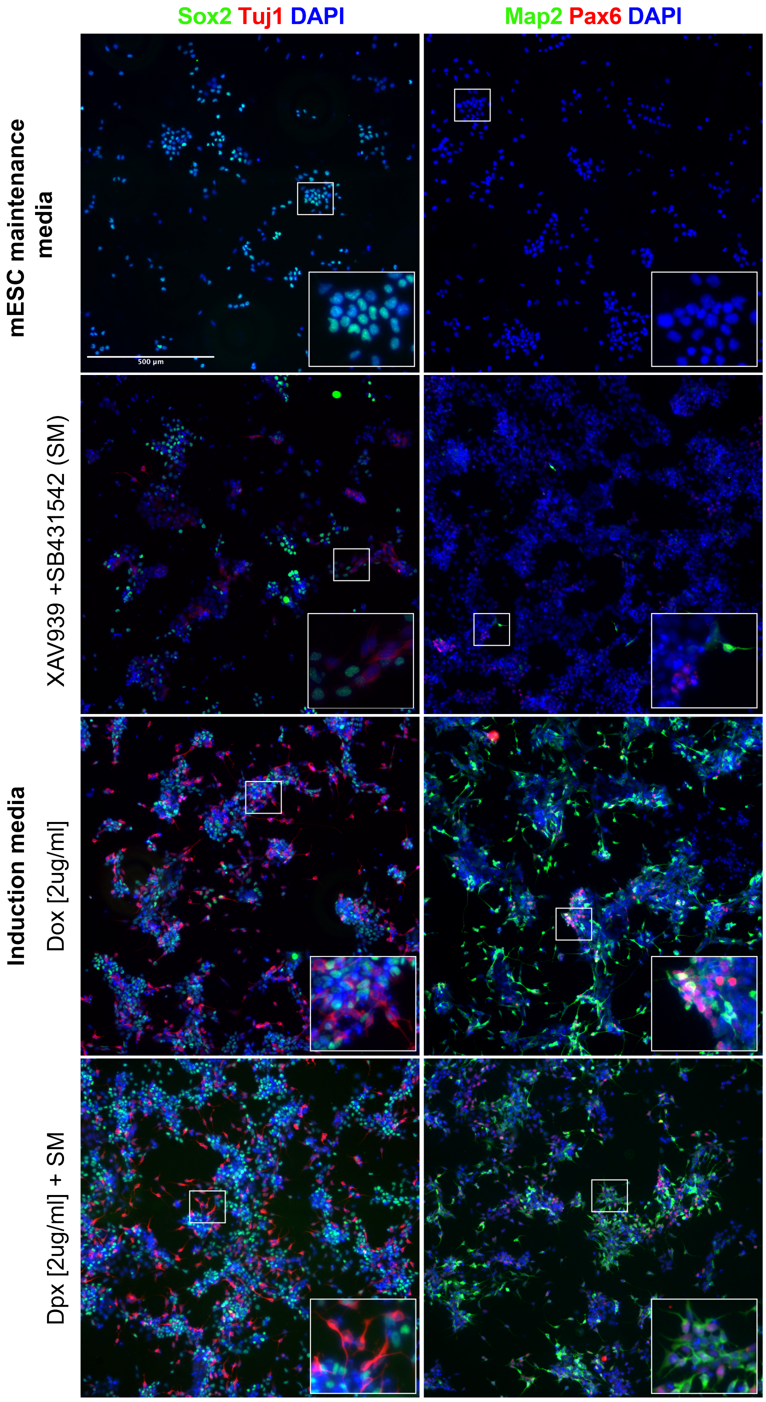


**Figure S2. Comparison of four culture protocols for early neural induction from mESCs.** Related to figure 1. Representative immunofluorescence images comparing four differentiation conditions at early time points. Columns display Sox2/Tuj1/DAPI staining (left) and Map2/Pax6/DAPI staining (right). Differentiation conditions are presented from top to bottom as follows: mESC maintenance media (control), showing limited neural differentiation; induction media supplemented with XAV939 and SB431542, promoting early neural patterning; induction media containing doxycycline (2 $\mu$g/mL) to induce *NGN2* expression; and induction media combining XAV939/SB431542 with doxycycline, resulting in enhanced neuronal differentiation. Sox2 marks neural progenitors, Tuj1 and Map2 label neuronal populations, Pax6 indicates early neural identity, and nuclei are counterstained with DAPI.

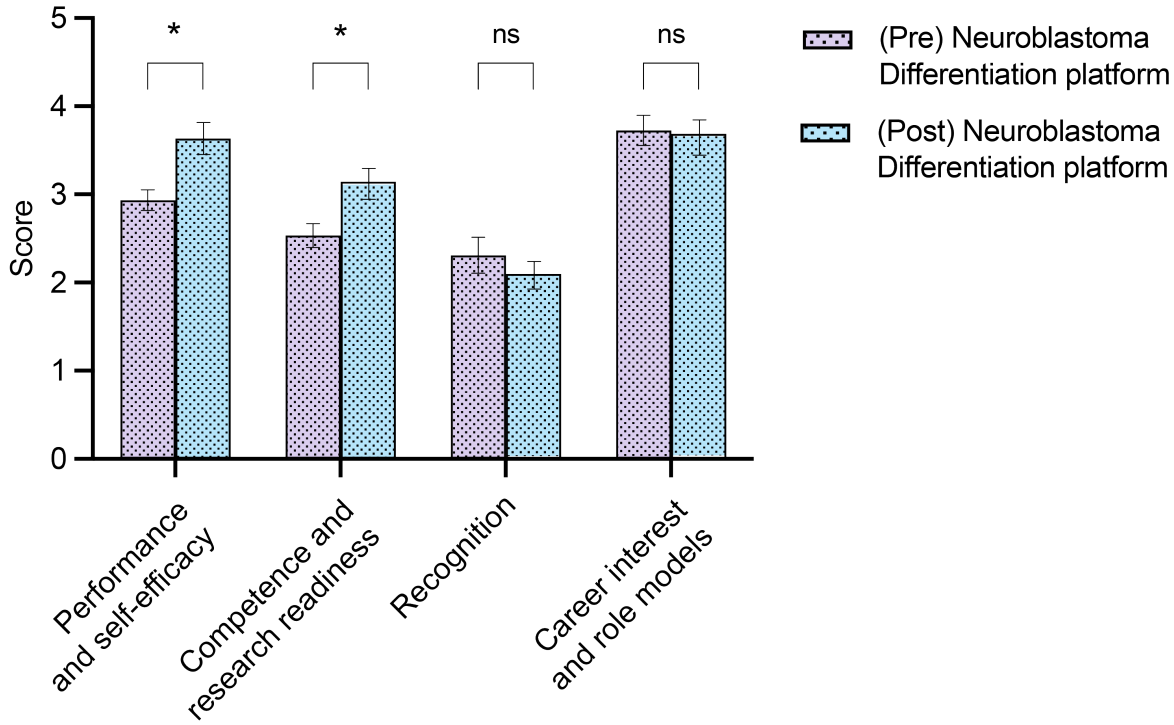


**Figure S3. Comparison of pre- and post-intervention scores across identity dimension on Neuroblastoma Differentiation Platform cohort at Alisal High School.** Related to figure 4. $n = 21$ Bars represent mean ± SEM. Unpaired t-test was performed across groups. *$p < 0.05$; **$p < 0.01$; ***$p < 0.001$.

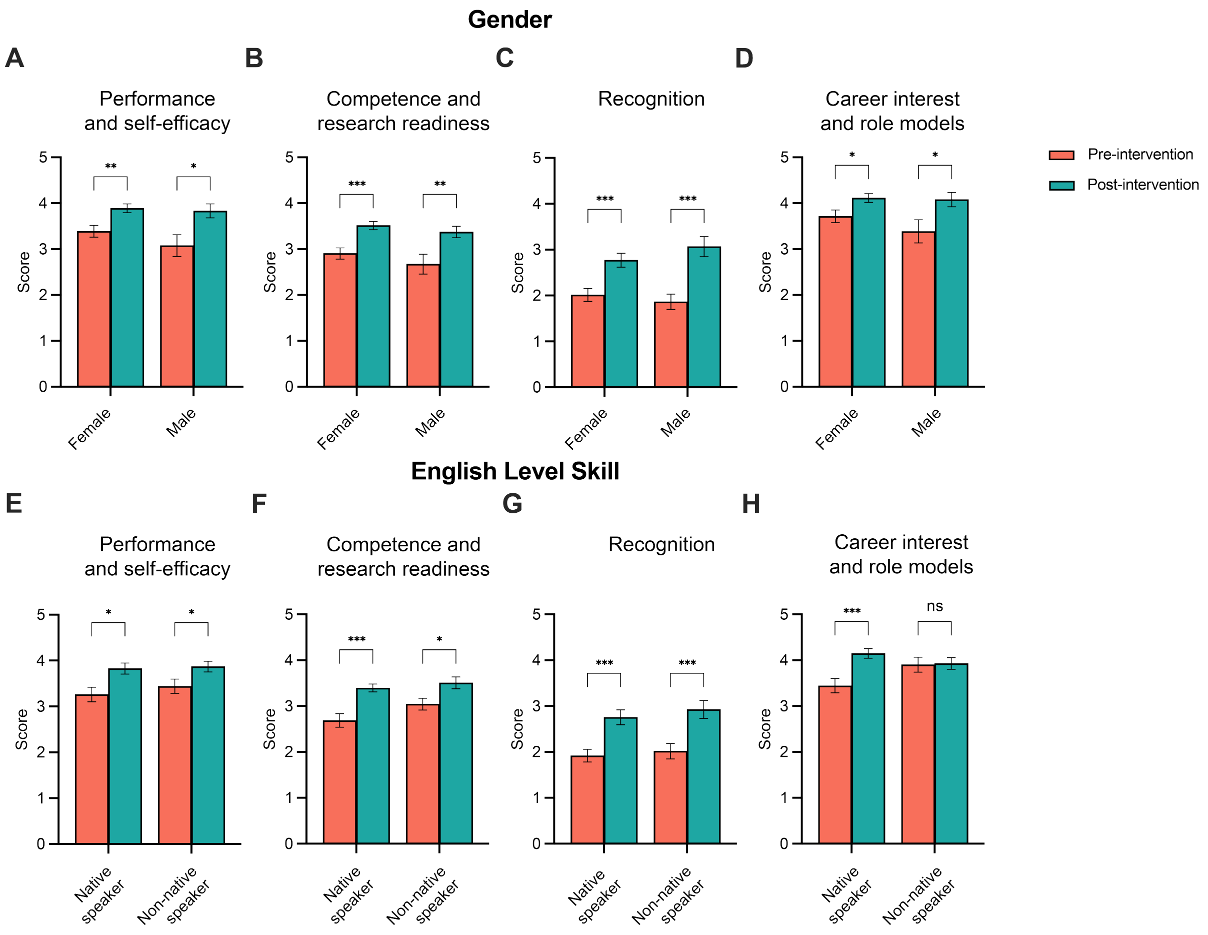


**Figure S4. Identity dimension scores stratified by gender and English proficiency.** Related to figure 4. Gender and English level skills across identity dimension scores using aggregated *NGN2* differentiation platform cohorts from Alisal High School and Berkeley City College. (**a–d**) Gender: female ($n = 48$) and male ($n = 22$). (**e–h**) English proficiency: native speakers ($n = 46$) and non-native speakers ($n = 28$). Bars represent mean ± SEM. Group differences were assessed using unpaired two-tailed t-tests. *$p < 0.05$; **$p < 0.01$; ***$p < 0.001$.

**Table S1.** **List of items in the Identity Questionnaire**. Related to Figure 2

| Scale | Item |
|---|---|
| **Science Performance Scale** | |
| P1 | I believe I perform well in my stem cell biology classes |
| P2 | I am able to achieve a good grade in my stem cell biology classes |
| P3 | I can complete my stem cell biology homework assignments |
| P4 | I am skilled in using tools and operating apparatus in stem cell experiments |
| P5 | I can conduct research in stem cell biology |
| **Science Competence Scale** | |
| C1 | I think I am good at stem cell biology |
| C2 | I can use stem cell biology concepts to explain natural phenomena |
| C3 | I believe I have a strong understanding of stem cell biology |
| C4 | I am confident explaining key concepts in stem cell biology |
| C5 | I believe I can understand even the most challenging aspects of stem cell biology with effort and dedication |
| **Science Recognition Scale** | |
| R1 | I consider myself a stem cell scientist |
| R2 | My classmates recognize me as a stem cell scientist |
| R3 | My teachers recognize me as a stem cell scientist |
| R4 | My family and friends recognize me as a stem cell scientist |
| **Science Interest Scale** | |
| I1 | I will continue to expand my knowledge of stem cell biology |
| I2 | I enjoy participating in stem cell biology-related activities |
| I3 | I believe the stem cell biology knowledge taught in my classes is valuable in the real world |
| I4 | I enjoy attending classes related to stem cell biology |
| I5 | I am planning to pursue a career related to stem cell biology |
| I6 | I would feel comfortable talking to people in stem cell biology-related careers |
| **Role Models Scale** | |
| RM1 | I believe having stem cell scientist role models will increase my interest in stem cell research |
| RM2 | I believe stem cell scientist role models are crucial for encouraging me to pursue stem cell careers |
| RM3 | Learning about the achievements of stem cell scientists will boost my confidence |
| RM4 | I like learning more about the achievements of stem cell scientists |

**Table S2.** **Demographics of Students Enrolled in BME80G - Bioethics in the 21st Century: Science, Business, and Society. University of California Santa Cruz. (N = 133).** Related to Figure 2.

| Category | Group | Count | % |
|---|---|---|---|
| Gender | Female | 71 | 53.4 |
| | Male | 55 | 41.4 |
| | Non-binary | 4 | 3.0 |
| | Prefer not to say | 2 | 1.5 |
| | Queer | 1 | 0.8 |
| Race / Ethnicity | Hispanic/Latinx or Spanish Origin | 17 | 12.8 |
| | White / European | 27 | 20.3 |
| | Asian / Asian-American | 60 | 45.1 |
| | Black / African American | 1 | 0.8 |
| | Middle Eastern / North African | 3 | 2.3 |
| | Native Hawaiian / Pacific Islander | 1 | 0.8 |
| | Jewish | 1 | 0.8 |
| | Bi-racial / Multi-racial | 22 | 16.5 |
| | Slavic | 1 | 0.8 |
| Language | Native speaker | 98 | 73.7 |
| | Non-native speaker | 35 | 26.3 |
| Parental Education | No parental education | 34 | 25.6 |
| | 4-year degree or higher | 99 | 74.4 |

**Table S3.** **Student Demographics for Alisal High School 2024 Cohort (N = 21).** Related to Figures 2 and 4.

| Category | Group | Count | % |
|---|---|---|---|
| Gender | Female | 16 | 76.2 |
| | Male | 5 | 23.8 |
| Race / Ethnicity | Hispanic/Latinx or Spanish Origin | 21 | 100 |
| Language | Native speaker | 12 | 57.1 |
| | Non-native speaker | 9 | 42.9 |
| Parental Education | No parental education | 17 | 81 |
| | 4-year degree or higher | 4 | 19 |

**Table S4.** **Student Demographics for Alisal High School 2025 Cohort (N = 32).** Related to Figures 2 and 4.

| Category | Group | Count | % |
|---|---|---|---|
| Gender | Female | 25 | 78.1 |
| | Male | 7 | 21.9 |
| Race / Ethnicity | Asian / Asian-American | 3 | 9.4 |
| | Bi-racial / Multi-racial | 4 | 12.5 |
| | Hispanic/Latinx or Spanish Origin | 25 | 78.1 |
| Language | Native speaker | 16 | 50 |
| | Non-native speaker | 16 | 50 |
| Parental Education | No parental education | 26 | 81.3 |
| | 4-year degree or higher | 6 | 18.8 |

**Table S5.** **Student Demographics for Berkeley City College 2023 Cohort (N = 14).** Related to Figures 2 and 4.

| Category | Group | Count | % |
|---|---|---|---|
| Gender | Female | 6 | 42.9 |
| | Male | 4 | 28.6 |
| | Non-binary | 3 | 21.4 |
| | Prefer not to say | 1 | 7.1 |
| Race / Ethnicity | Asian / Asian-American | 7 | 50.0 |
| | Black / African American | 1 | 7.1 |
| | Hispanic/Latinx or Spanish Origin | 2 | 14.3 |
| | Middle Eastern / North African | 1 | 7.1 |
| | White / European | 3 | 21.4 |
| Language | Native speaker | 10 | 71.4 |
| | Non-native speaker | 4 | 28.6 |
| Parental Education | No parental education | 6 | 42.9 |
| | 4-year degree or higher | 8 | 57.1 |

**Table S6.** **Student Demographics for Berkeley City College 2024 Cohort (N = 16).** Related to Figures 2 and 4.

| Category | Group | Count | % |
|---|---|---|---|
| Gender | Female | 10 | 62.5 |
| | Male | 6 | 37.5 |
| Race / Ethnicity | Asian / Asian-American | 3 | 18.8 |
| | Black / African American | 1 | 6.3 |
| | Hispanic/Latinx or Spanish Origin | 6 | 37.5 |
| | White / European | 6 | 37.5 |
| Language | Native speaker | 10 | 62.5 |
| | Non-native speaker | 6 | 37.5 |
| Parental Education | No parental education | 7 | 43.8 |
| | 4-year degree or higher | 9 | 56.3 |

**Table S7.** **Student Demographics for Berkeley City College 2025 Cohort (N = 12).** Related to Figures 2 and 4.

| Category | Group | Count | % |
|---|---|---|---|
| Gender | Female | 7 | 58.3 |
| | Male | 5 | 41.7 |
| Race / Ethnicity | Asian / Asian-American | 2 | 16.7 |
| | Bi-racial / Multi-racial | 4 | 33.3 |
| | Black / African American | 1 | 8.3 |
| | Hispanic/Latinx or Spanish Origin | 1 | 8.3 |
| | Middle Eastern / North African | 1 | 8.3 |
| | White / European | 3 | 25.0 |
| Language | Native speaker | 10 | 83.3 |
| | Non-native speaker | 2 | 16.7 |
| Parental Education | No parental education | 3 | 25 |
| | 4-year degree or higher | 9 | 75 |

**Table S8.** **Final Identity Questionnaire items following confirmatory factor analysis (CFA) - Stem Cell Research Identity Scale (SCRIS) Instrument**. Related to Figure 2.

| Scale | Item |
|---|---|
| **Perform and Self-efficacy Scale** | |
| P1 | I believe I perform well in my stem cell biology classes |
| P2 | I am able to achieve a good grade in my stem cell biology classes |
| P3 | I can complete my stem cell biology homework assignments |
| **Competence and Research Readiness Scale** | |
| P5 | I can conduct research in stem cell biology |
| C1 | I think I am good at stem cell biology |
| C2 | I can use stem cell biology concepts to explain natural phenomena |
| C3 | I believe I have a strong understanding of stem cell biology |
| C5 | I believe I can understand even the most challenging aspects of stem cell biology with effort and dedication |
| **Recognition Scale** | |
| R1 | I consider myself a stem cell scientist |
| R2 | My classmates recognize me as a stem cell scientist |
| R3 | My teachers recognize me as a stem cell scientist |
| R4 | My family and friends recognize me as a stem cell scientist |
| **Career Interest and Role Model Impact Scale** | |
| I4 | I enjoy attending classes related to stem cell biology |
| I5 | I am planning to pursue a career related to stem cell biology |
| RM1 | I believe having stem cell scientist role models will increase my interest in stem cell research |
| RM3 | Learning about the achievements of stem cell scientists will boost my confidence |

#### Supplementary Note 1. Microscope Assembly Overview

This supplementary section provides detailed assembly instructions, component specifications, and software configuration for the distributed cloud-connected microscope used in this study. The overall system architecture, imaging workflow, and experimental operating conditions are described in the Methods section under *Distributed Cloud-Connected Microscopy Architecture*.

Complete hardware documentation, control software, and streaming scripts for this microscope implementation are available in the accompanying GitHub repository:

**https://github.com/braingeneers/educationpaper_supplemental_microscope**

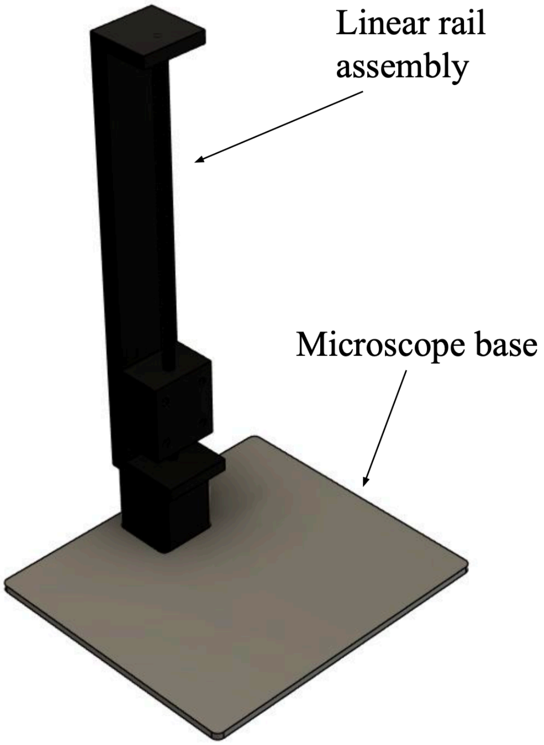


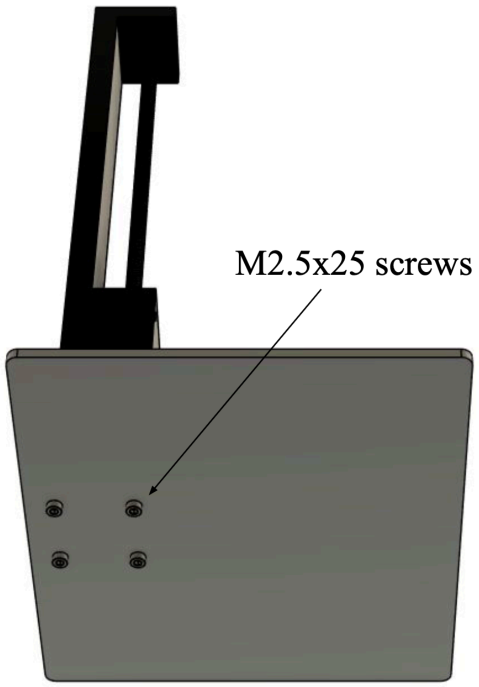


Remove the four M2.5 screws from the bottom of the linear rail assembly and replace them with four M2.5x25 screws, using them to secure the base of the microscope to the linear rail.

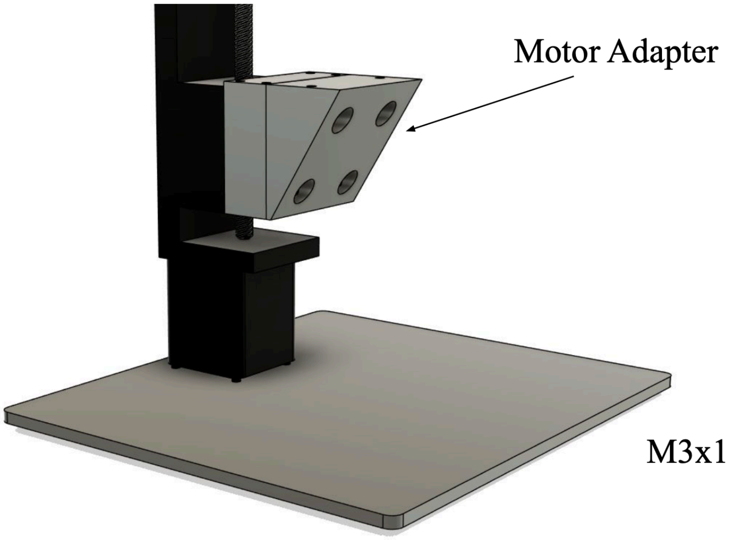

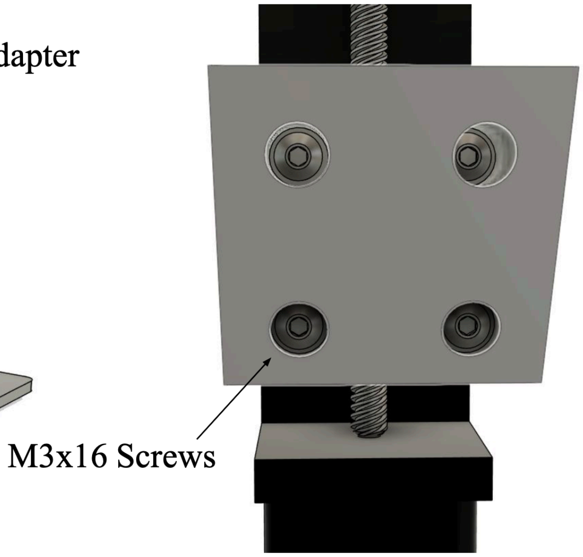


Mount the motor adapter to the linear rail using four M3x16 screws.

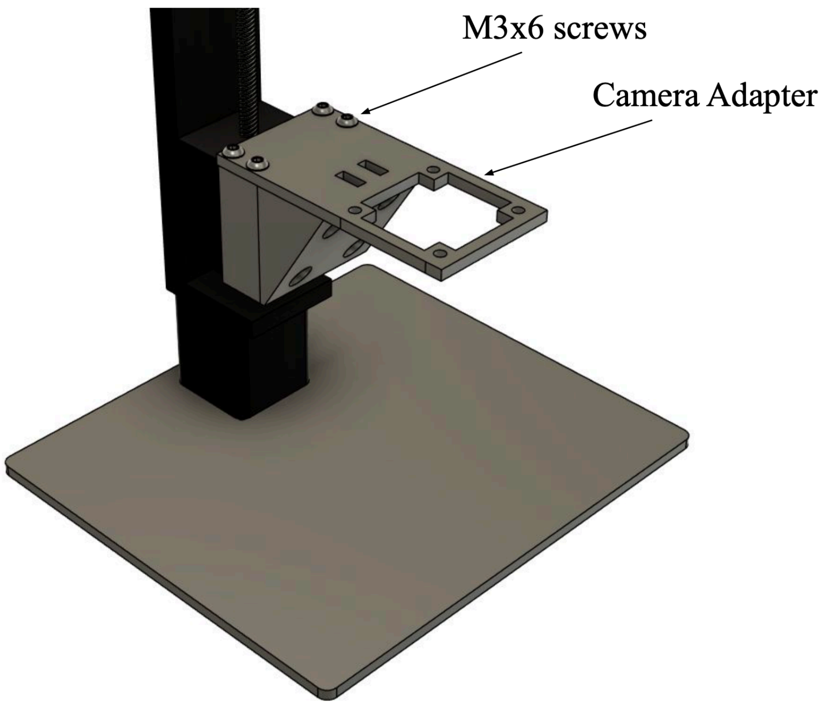


Mount the camera adapter to the motor adapter using four M3x6 screws.

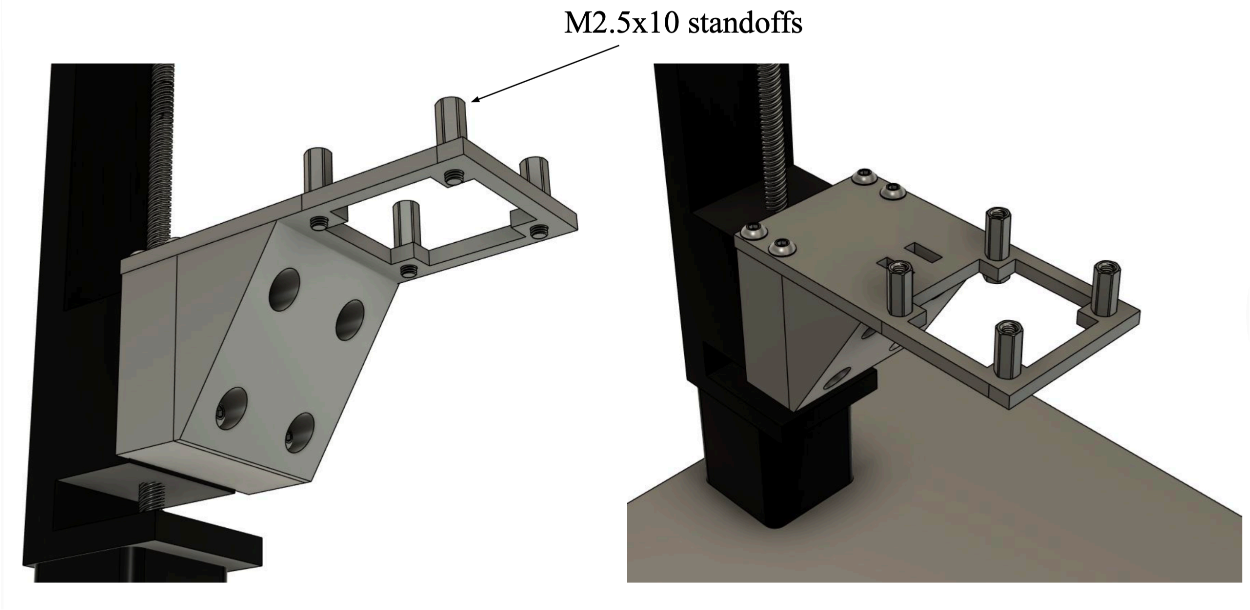


Mount four M2.5x10 standoffs to the holes on the end of the camera adapter.

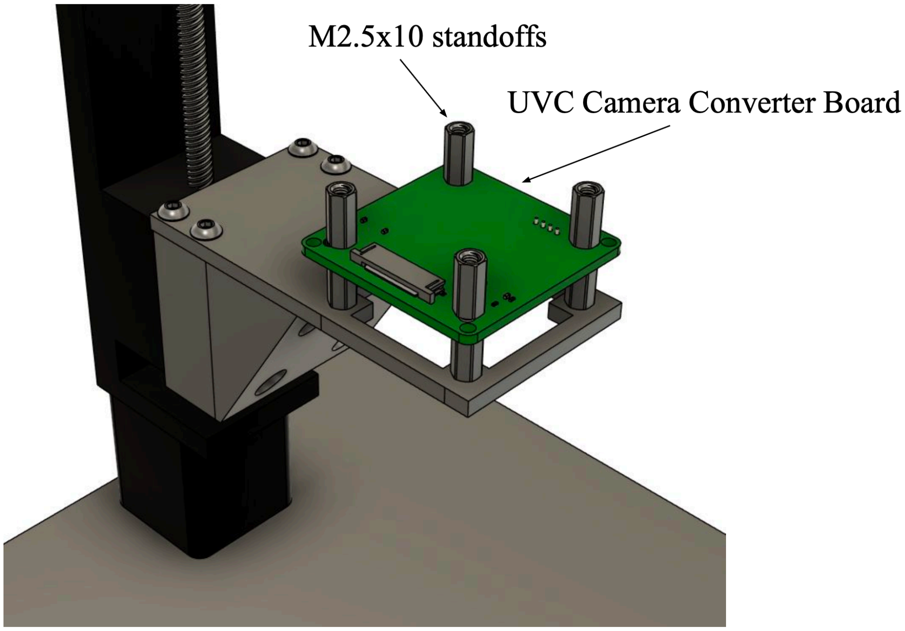


Secure the UVC Camera Converter Board using four more M2.5x10 standoffs.

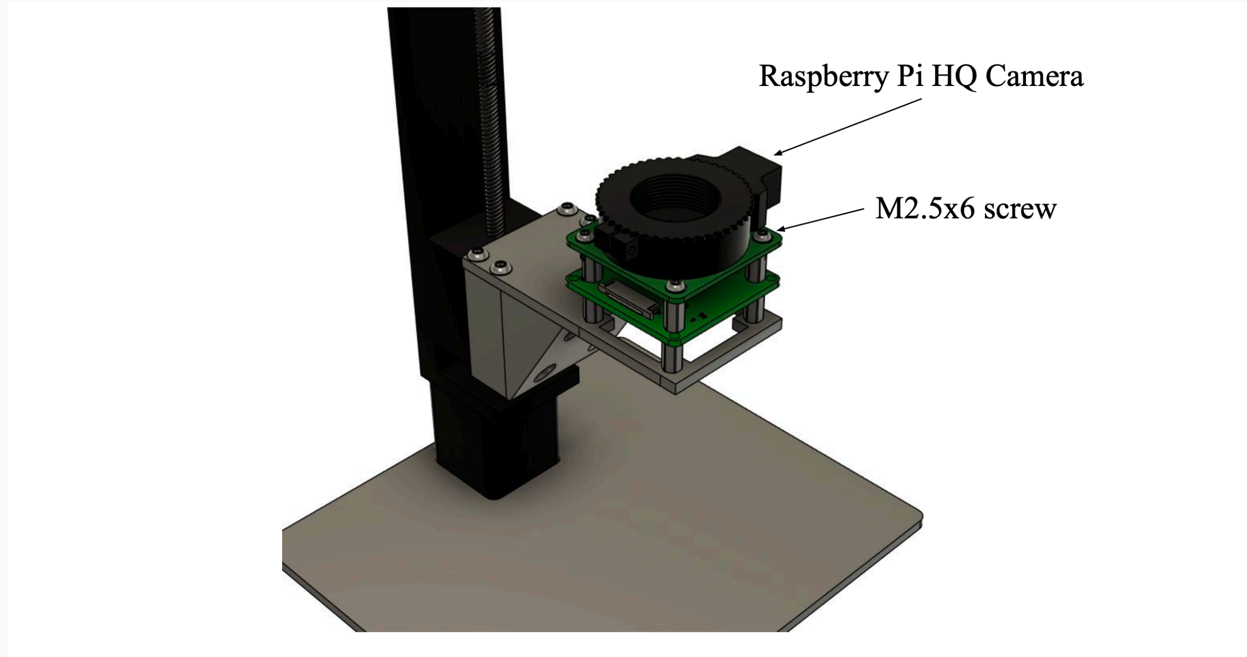


Place the Raspberry Pi HQ Camera on top of the upper set of standoffs and secure it using four M2.5x6 screws.

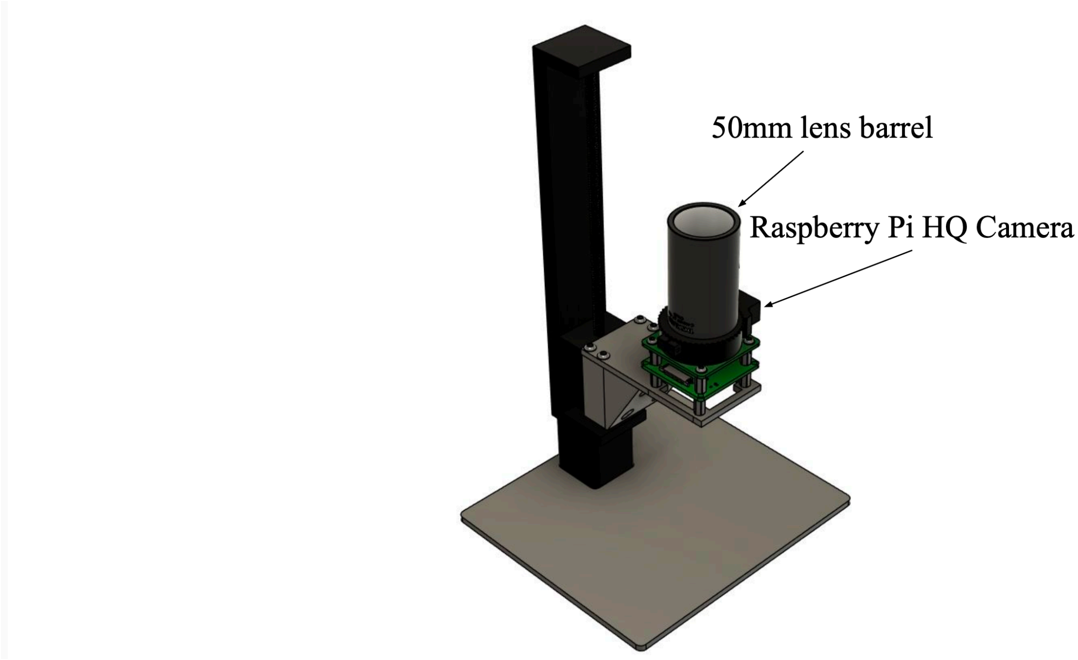


Mount the 50mm lens barrel on top to the Raspberry Pi HQ Camera.

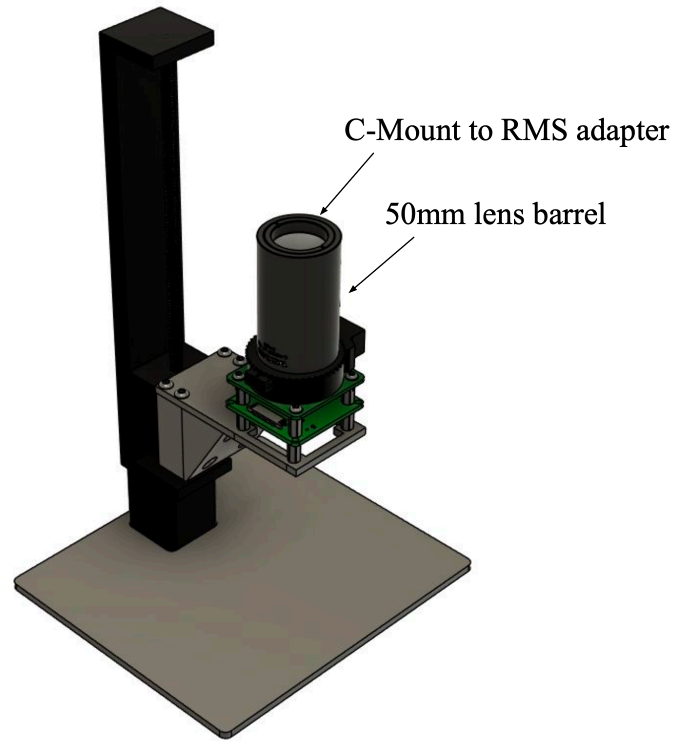


Seat the C-Mount to RMS adapter at the top of the lens barrel.

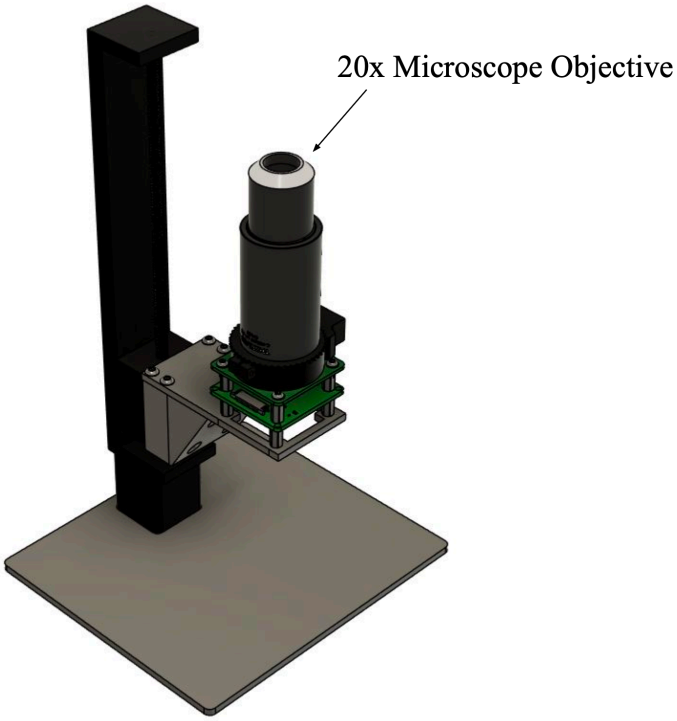


Secure the 20x microscope objective into the C-Mount to RMS adapter.

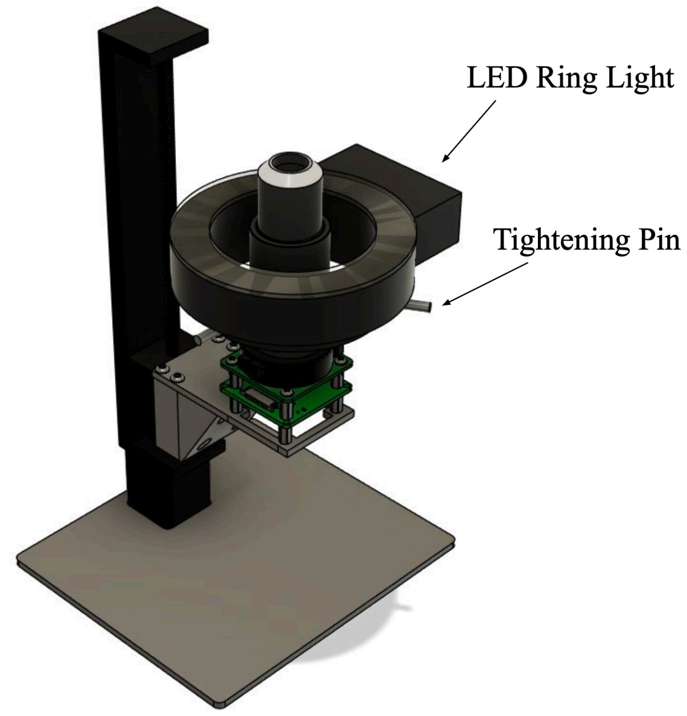


Secure the ring light to the lens barrel using the three tightening pins.

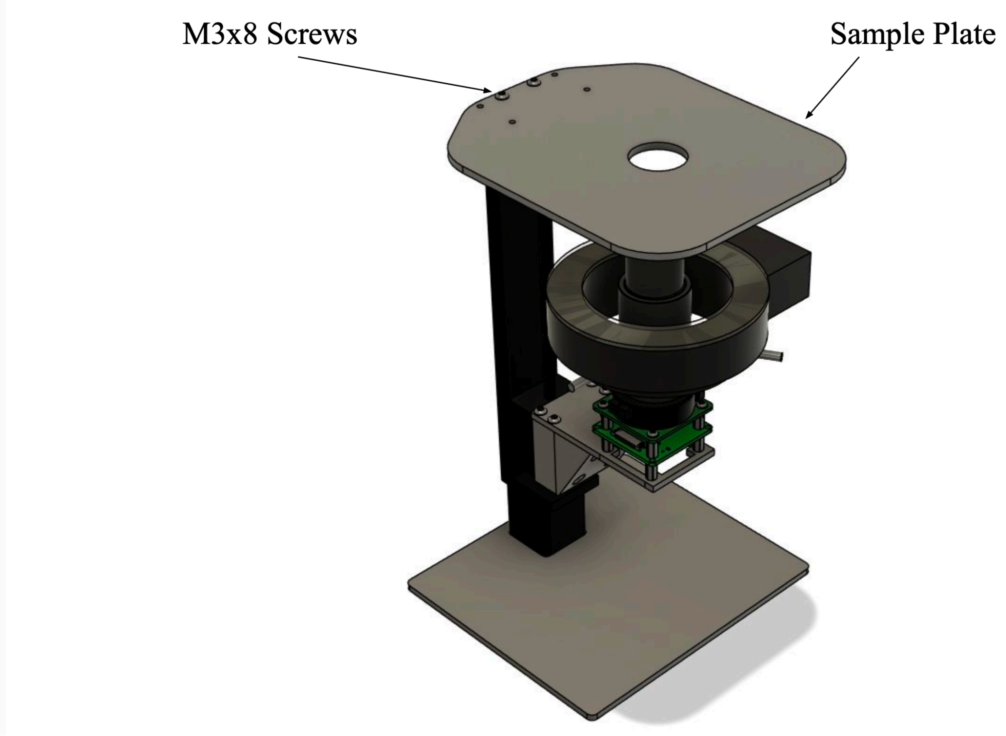


Remove the two M3 screws at the top of the linear rail assembly and replace them with M3x8 screws, using them to secure the sample plate in place.
